## Supporting Informations for "Bridging the gap between the evolutionary dynamics and the molecular mechanisms of meiosis : a model based exploration of the *PRDM9* intra-genomic Red Queen"

#### Supplementary Appendix S1 : Derivation of the analytical approximation to the equilibrium regime of the Red Queen

### 1 Introduction

In this document, we give the detailed derivation of the analytical approximation for the stationary regime of the Red Queen process. The derivation proceeds along similar lines as in Latrille *et al.* [1]. It relies on a self-consistent mean field argument. In brief, the derivation assumes the following hypotheses :

- Wright-Fisher model (*i.e.* non-overlapping generations) with mutation and selection
- Panmictic population (random mating)
- Constant population size $N$
- Highly polymorphic model
- Weak erosion
- Strong selection implying that genetic drift can be ignored

The first four hypotheses are entailed by the simulator. The last three are additional assumptions that are made in order to make the analytical derivation feasible. The analytical results will therefore be valid only in regimes in which these three assumptions are met.

Based on these hypotheses, we first express the frequency $f(t)$ and the proportion of active site , $\theta(t)$ through time of a typical *PRDM9* allele. We will then obtain expressions for the erosion level of an allele as a function of its age, and the mean erosion level of the population. Relying on the argument that the population itself is composed of such typical *PRDM9* alleles successively invading, with a mean time interval between successive invasions noted $\tau$, itself depending on the mean fitness of the population, we can obtain a self-consistent expression for the mean erosion level, from where we can then express all summary statistics of interest.

In terms of notations, since we consider that all alleles follow the same trajectory over their existence, differing only in their arrival time in the population, all quantities will be expressed as a function of $t$ (e.g. $f(t)$ for the frequency, or $\theta(t)$ for the proportion of active sites for the allele),

$t$ being understood as the time since the birth of the allele. This thus differs from the notations in the main text, where time is absolute and measured relative to a single origin (defined as the time when the simulation run has reached equilibrium, at which point the summary statistics start to be monitored), and quantities are explicitly indexed by the specific allele or allelic combination for which they are being computed (e.g. $f_{i,t}$ for the frequency of allele $i$ at time $t$, or $w^{het}_{i,j,t}$ for the fitness of an individual with alleles $i$ and $j$ at time $t$).

In addition, below, we will consider a change of variable, expressing the same quantities as a function, not of time $t$, but as a function of the allele intrinsic age $z$ (which is defined below and which is an increasing function of $t$). With a slight abuse of notations, the same symbol will be used as a function of $t$ or $z$ (e.g. $f(t)$ or $f(z)$, or even simply $f$), given that the meaning will be clear depending on the context.

### 2 Analytical developments

#### 2.1 Studied model

First, assuming negligible random drift, the evolution of the frequency through time $f(t)$ of a typical allele is deterministic and is given by:

$$\frac{df}{dt} = \frac{w^*(t) - \bar{w}}{\bar{w}} f, \tag{1}$$

where $w^*(t)$ is the mean fertility of the allele at time $t$ and $\bar{w}$ is the mean fertility of all *PRDM9* alleles in the population (weighted by their frequency). Of note, under the mean-field approximation considered here, $\bar{w}$ is, by definition, assumed independent of time. Furthermore, it is considered as an independent variable, which will be determined in a second step, based on a self-consistent argument.

Equation 1 means that selection will depend on differential erosion. In turn, erosion depends on the frequency trajectory of the allele. To decouple these effects, we start by observing that the fraction of active target sites for the allele varies approximately as :

$$\frac{d\theta}{dt} \approx -\rho f \theta \tag{2}$$

In this equation, $\rho$ is the erosion rate per generation. It can be expressed as :

$$\rho = (2Nv)(2g) \tag{3}$$

where $g = \frac{d}{8h}$ is the mean gene conversion rate (*i.e.* the probability that a site undergoes a DSB and a repair). Thus, $\rho$ is equal to :

$$\rho = \frac{Nvd}{2h} \tag{4}$$

Equation (2) expresses the fact that the rate of extinction of target sites is equal to the mutation rate at the level of the population ($2Nv$) multiplicated by the probability of fixation of the inactive mutant. Assuming strong gene conversion ($4Ng >> 1$), this probability is well approximated by twice the conversion rate, *i.e.* $2fg$. Note that the rate of erosion given by (3) is only approximate. A more accurate expression accounting for the affinity distribution of the sites will be given below.

#### 2.2 The intrinsic age of an allele ($z$)

Therefore,

$$z(t) = \rho \int_0^t f(u)\, du \tag{5}$$

can be seen as a measure of the cumulative erosion level of an allele. As such, it can be used as a measure of the intrinsic age of the allele. We can then express the equation describing the evolution of $f$ and $\theta$ directly as a function of $z$, instead of $t$. This change of variable will entail the following factor :

$$\frac{dz}{dt} = \rho f. \tag{6}$$

As mentioned above, our derivation assumes weak erosion. Mathematically, this translates into the assumption that $z << 1$.

#### 2.3 Frequency of an allele (f) depending on its age (z) : f(z)

Then, we want to express the evolution of the frequency of an allele in the population as a function on its intrinsic age $z$:

$$\frac{df}{dz} = \frac{df}{dt}\frac{dt}{dz} = \left[\frac{w^*(z) - \bar{w}}{\bar{w}}\right] \cdot \left[\frac{1}{\rho}\right] = \frac{1}{\rho}\left(\frac{w^*(z)}{\bar{w}} - 1\right) \tag{7}$$

However, $\frac{w^*}{\bar{w}}$ is extremely close to 1, so $\frac{w^*}{\bar{w}} - 1$ is close to 0. In addition, we know that, in the vicinity of 0, $x \approx ln(1 + x)$. As a result, we can write

$$\frac{1}{\rho}\left(\frac{w^*(z)}{\bar{w}} - 1\right) \approx \frac{1}{\rho} ln \left(1 + \left(\frac{w^*(z)}{\bar{w}} - 1\right)\right) = \frac{1}{\rho} ln \left(\frac{w^*(z)}{\bar{w}}\right) \tag{8}$$

Working in the weak erosion limit ($z << 1$) allows us to linearize $ln(w)$ in the vicinity of 0 :

$$ln\left(\frac{w^*(z)}{\bar{w}}\right) = ln(w^*(z)) - ln(\bar{w}) \approx ln(w(0,0)) - \frac{\alpha}{2}(z + \bar{z}) - ln(w(0,0)) + \frac{\alpha}{2}(\bar{z} + \bar{z}) = -\frac{\alpha}{2}(z - \bar{z}) \tag{9}$$

where:

$$\alpha = \left|\frac{\partial ln(w)}{\partial z}\right|_{(z=0)} \tag{10}$$

is the slope at the origin of $ln(w)$. It depends on the mechanistic parameters, in a way that will be determined further below.

Then, by replacement in equation (7) :

$$\boxed{\frac{df}{dz} \approx -\frac{\alpha}{2\rho}(z - \bar{z})} \tag{11}$$

Integrating equation (11), with the constraint that $f(0) = 0$, gives :

$$f(z) = -\frac{\alpha}{4\rho} z[z - 2\bar{z}]. \tag{12}$$

The function $f$ has the shape of a concave parabola and $f(z) = 0$ when $z = 0$ and $z = 2\bar{z}$.

##### 2.4 Mean age of an allele in the population ($\bar{z}$) : a self-consistent derivation

Equations (11) and (12) depend on $\bar{z}$, and thus we now need to express $\bar{z}$ as a function of the model parameters.
The mean allele age of the population $\bar{z}$ is equal to :

$$\bar{z} = \sum_i (f_i z_i). \tag{13}$$

where, for simplicity, we have momentarily used explicit indexing of multiple alleles simultaneously segregating in the population at a given time.

Relying on a tiling argument (Latrille *et al.*, 2017 [1]) this can also be expressed as :

$$\bar{z} = \frac{1}{\tau} \int_0^{+\infty} f(t) z(t) \, dt. \tag{14}$$

Equation (14) expresses the idea that, at stationarity, the allele frequency distribution at a given time point (eq. (13)) is equivalent to the distribution of frequencies at which a typical allele has segregated over its entire life normalized by the mean waiting time $\tau$ between successive invasions (see Latrille *et al.*, 2017 figure 6 [1]).
We then do a change of variable from $t$ to $z$ in the integrand:

$$\bar{z} = \frac{1}{\tau} \int_0^{z(\infty)} z f \frac{dt}{dz} \, dz = \frac{1}{\tau} \int_0^{z(\infty)} z f \left( \frac{dz}{dt} \right)^{-1} dz, \tag{15}$$

then, we replace $\frac{dz}{dt}$ by its expression in equation (2) :

$$\bar{z} = \frac{1}{\tau} \int_0^{z(\infty)} z f \frac{1}{f\rho} \, dz = \frac{1}{\tau\rho} \int_0^{z(\infty)} z \, dz = \frac{1}{\tau\rho} \frac{z(\infty)^2}{2}. \tag{16}$$

In order to determine $z(\infty)$, we express the constrain that $\sum_i (f_i) = 1$. This expression can also be written as :

$$1 = \sum_i (f_i) = \frac{1}{\tau} \int_0^{\infty} f(t) \, dt = \frac{1}{\tau} \int_0^{z(\infty)} f \frac{dt}{dz} \, dz = \frac{1}{\rho\tau} \int_0^{z(\infty)} f \frac{1}{f} \, dz = \frac{1}{\rho\tau} \int_0^{z(\infty)} dz = \frac{z(\infty)}{\rho\tau} \tag{17}$$

So

$$z(\infty) = \rho\tau, \tag{18}$$

such that, by replacement :

$$\bar{z} = \frac{\rho\tau^2}{2\rho\tau} = \frac{\rho\tau}{2}. \tag{19}$$

We now need to express $\tau$, which is the inverse of the invasion rate of a new allele in the population. The rate of invasion is equal to the rate of mutation at the population level ($2Nu$) multiplicated by the invasion probability. Assuming strong selection, this probability is well approximated by $2s_0$.

$$\tau^{-1} = (2Nu).(2s_0) \tag{20}$$

where $u$ is the mutation rate at the Prdm9 locus and $s_0$ is the selection coefficient of a new allele in the population. Based on equation (9), $s_0$ can be expressed as $s_0 = \frac{\alpha}{2}\bar{z}$. Thus :

$$\tau^{-1} = 4Nu\frac{\alpha}{2}\bar{z} = \mu\frac{\alpha}{2}\bar{z}, \tag{21}$$

Where $\mu = 4Nu$. If we replace this in equation (19) we finally obtain :

$$\bar{z} = \frac{\rho}{2\tau^{-1}} = \frac{\rho}{2}\frac{2}{\mu\alpha\bar{z}} = \frac{\rho}{\mu\alpha\bar{z}} \iff \bar{z}^2 = \frac{\rho}{\mu\alpha} \iff \boxed{\bar{z} = \sqrt{\frac{\rho}{\mu\alpha}}} \tag{22}$$

We thus recover the main result of the derivation given in Latrille *et al.* [1], although now, the compound parameters $\rho$ and $\alpha$ depend on the mecanistic details of our model. We have already seen that $\rho = \frac{Nvd}{2h}$. On the other hand, we still need to express $\alpha$.

#### 2.5 Slope at the origin of the fertility rate : $\alpha$

As mentioned above, $\alpha$ is the slope of $ln(w)$ at the origin (*i.e.* for two new alleles of age $z_1 = z_2 = 0$):

$$\alpha = \left|\frac{\partial \ln(w(z_1,z_2))}{\partial z_1}\right|_{(z_1=0,z_2=0)} = \left|\frac{1}{w}\frac{\partial w(z_1,z_2)}{\partial z_1}\right|_{(z_1=0,z_2=0)} = \left|\frac{\partial w(z_1,z_2)}{\partial z_1}\right|_{(z_1=0,z_2=0)}, \tag{23}$$

since $w \approx 1$.
Thus, in order to find an explicit expression for $\alpha$, we need to express $w$ according to the parameters.
Assuming that gametes are not limiting, the fitness of an individual is equal to the rate of success of meiosis, which is itself equal to the probability of having at least one DSB in a symmetrical bound site. So, we can write $w$ as 1 - the probability of having no DSB in symmetrical bound site (1 - probability of failure of the meiosis). The number of DSBs in symmetrical bound sites is approximately Poisson of mean $dq(z_1, z_2)$, where $d$ is the mean number of DSBs and $q(z_1, z_2)$ is the probability that a DSB occurs in a symmetrically bound site in an individual heterozygous for two *PRDM9* alleles of age $z_1$ and $z_2$. Thus, the probability of 0 DSB in a symmetrically bound site is $e^{-dq(z_1,z_2)}$. So, $w(z_1, z_2)$ can be expressed as :

$$w(z_1,z_2) = 1 - e^{-dq(z_1,z_2)} \tag{24}$$

Substituting in equation (23), we obtain :

$$\alpha = \left|\frac{\partial w(z_1,z_2)}{\partial z_1}\right|_{(z_1=0,z_2=0)} = \left| d\,\frac{\partial q(z_1,z_2)}{\partial z_1}\right|_{(z_1=0,z_2=0)} e^{-dq(z_1=0,z_2=0)}\Bigg| \tag{25}$$

We see here that, in order to obtain an explicit formula for $\alpha$, it is necessary to express $q(z_1, z_2)$ as a function of the model parameters and then compute its derivative as a function of $z$.

#### 2.6 Probability of symmetrical binding : q

For a given site of affinity $y$, the probability that PRDM9 is bound is given by $x = \frac{cy}{1+cy}$ where $c = 1$ or $c = 2$. The conditional probability of symmetrical binding at that site (conditional on at least one of the four chromatids being bound) is then equal to :

$$q = \frac{2x^2 - x^3}{x} \tag{26}$$

This expression is valid for a single site. To compute the mean probability of symmetrical binding over the genome, we need to average the numerator and the denominator separately over the affinity distribution across sites. Of note this distribution itself depends on the age $z$ of the allele, and the mean over the distribution of a given function $B(y)$ is noted $< B >_z$. Also, the mean q over the genome depends on the *Prdm9* genotype of this individual. In the case of a homozygote, thus possessing twice the same *Prdm9* allele of age $z$, $q$ is equal to:

$$q^{hom}(z) = \frac{2 < x^2 >_z - < x^3 >_z}{< x >_z} \tag{27}$$

If the individual is heterozygous for *Prdm9* with two alleles of age $z_1$ and $z_2$, $q$ is equal to:

$$q^{het}(z_1, z_2) = \frac{2 < x^2 >_{z_1} - < x^3 >_{z_1} + 2 < x^2 >_{z_2} - < x^3 >_{z_2}}{< x >_{z_1} + < x >_{z_2}} \tag{28}$$

The mean over the affinity distribution can be more precisely expressed as :

$$< B >_z = \int B(y)\theta_y(z)\varphi(y)\, dy. \tag{29}$$

In this equation $\varphi(y)$ is the affinity distribution of an allele at birth and $\theta_y(z)$ is the fraction of active sites with a given affinity $y$ recognised by an allele of age $z$. Thus, $\theta_y(z)\varphi(y)dy$ is the total number of target sites still active with an affinity $y \pm dy$.
The integrals of the form (29) can be obtained numerically. However, they depend on $\theta_y(z)$, which we therefore need to determine.

#### 2.7 Fraction of active sites with a given affinity $y$ recognised by an allele of an age $z$ : $\theta_y(z)$

Here our aim is to compute more precisely the proportion of target sites of a given affinity $y$ that are still active, for an allele of age $z$. We note this quantity $\theta_y(z)$

By an argument similar to that used for equation (3):

$$\frac{d\theta_y}{dt} = -(2Nv).(2fg_y^{het})\theta_y \tag{30}$$

where $g_y^{het} = g_y^{het}(z_1, z_2)$ is the gene conversion rate at sites of affinity $y$ in a genotype $(z_1, z_2)$. Of note, we consider only heterozygotes for *PRDM9* since we work under the assumption of a highly polymorphic regime. In turn :

$$g_y^{het} = d.\frac{\frac{cy}{1+cy}}{4h\left[\left\langle\frac{cy}{1+cy}\right\rangle_{z_1} + \left\langle\frac{cy}{1+cy}\right\rangle_{z_2}\right]} \tag{31}$$

The fraction on the right-hand side of equation (31) is the proportion of sites of affinity $y$ among all sites that are bound by either one of the two *PRDM9* alleles.

So, if we replace it in equation (30), we obtain :

$$\frac{d\theta_y}{dt} = -2Nv.d\frac{\frac{cy}{1+cy}}{4h\left[\left\langle\frac{cy}{1+cy}\right\rangle_{z_1} + \left\langle\frac{cy}{1+cy}\right\rangle_{z_2}\right]}.2f(t)\theta_y(z) = -Nv\frac{d}{h}.\frac{\frac{cy}{1+cy}}{\left[\left\langle\frac{cy}{1+cy}\right\rangle_{z_1} + \left\langle\frac{cy}{1+cy}\right\rangle_{z_2}\right]}.f(t)\theta_y(z). \tag{32}$$

If we suppose that $< \frac{cy}{1+cy} >_{z_1}$ and $< \frac{cy}{1+cy} >_{z_2}$ don't vary too much as a function of $z$ ($\approx < \frac{cy}{1+cy} >_0$), then we have:

$$\frac{d\theta_y}{dt} = -Nv\frac{d}{h}\frac{\frac{cy}{1+cy}}{2\left\langle \frac{cy}{1+cy} \right\rangle_0}.f(t)\theta_y(z) \tag{33}$$

Then, by setting $\rho = \frac{Nvd}{2h}$ we can re-express the equation (36) as

$$\frac{d\theta_y}{dt} = -\rho\frac{\frac{cy}{1+cy}}{\left\langle \frac{cy}{1+cy} \right\rangle_0}.f(t)\theta_y(z) \tag{34}$$

And at this stage we can pose $\gamma(y) = \frac{\frac{cy}{1+cy}}{\left\langle \frac{cy}{1+cy} \right\rangle_0}$, which gives the following equation:

$$\frac{d\theta_y}{dt} = -\rho f(t)\gamma(y)\theta_y. \tag{35}$$

Now, we can replace $\rho f(t)$ by $\frac{dz}{dt}$:

$$\frac{d\theta_y}{dt} = -\frac{dz}{dt}\gamma(y)\theta_y \Leftrightarrow \frac{d\theta_y(z)}{dz} = -\gamma(y)\theta_y \tag{36}$$

Finally if we integrate this equation, we obtain the fraction of active sites with an affinity $y$ recognized by an allele of age $z$ :

$$\theta_y(z) = \theta_y(0)e^{-\gamma(y)z} \tag{37}$$

By definition, $\theta_y(0)$ is the fraction of active sites with an affinity y recognized by an allele of age 0. Thus $\theta_y(0) = 1$, and as a result, equation (33) becomes

$$\boxed{\theta_y(z) = e^{-\gamma(y)z}} \tag{38}$$

Under the condition of weak erosion, we can simplify equation (38) which gives

$$\theta_y(z) \approx 1 - \gamma(y)z \tag{39}$$

and by averaging over the affinity, we get

$$< \theta_y(z) > \approx 1 - z \tag{40}$$

#### 2.8 Expression of $\alpha$

Thanks to this equation, we are now able to express the successive moments $< x^m >_z$ for $m = 1$, 2 and 3.

$$< x^m >_z = \int \left(\frac{cy}{1+cy}\right)^m \theta_y(z)\varphi(y)\,dy = \int \left(\frac{cy}{1+cy}\right)^m e^{-\gamma(y)z}\varphi(y)\,dy \tag{41}$$

To express $\alpha$, we need $q(0,0)$ and $\left.\frac{\partial q(z_1,z_2)}{\partial z_1}\right|_{z_1=0,z_2=0}$.
For $q(0,0)$ we have the following expression

$$q(0,0) = \frac{2 < x^2 >_0 - < x^3 >_0 + 2 < x^2 >_0 - < x^3 >_0}{< x >_0 + < x >_0} = \frac{2 < x^2 >_0 - < x^3 >_0}{< x >_0} \tag{42}$$

with $< B >_0 = \int B(y)\varphi(y)$.

And for $\left.\frac{\partial q(z_1,z_2)}{\partial z_1}\right|_{z_1=0,z_2=0}$, we first need to determine the general expression of $\frac{\partial <B>_z}{\partial z}$ and then express all the moments $< x^n >$ that we need.

$$\frac{\partial < B >_z}{\partial z} = \int B(y)\varphi(y)(-\gamma(y))e^{\gamma(y)z}\,dy = - < \gamma(y)B >_z \tag{43}$$

And with $\gamma(y) = \frac{\frac{cy}{1+cy}}{\left\langle \frac{cy}{1+cy}\right\rangle_0} = \frac{x}{<x>_0}$ we obtain :

$$\frac{\partial < B >_z}{\partial z} = -\frac{< xB >_z}{< x >_0} \tag{44}$$

So

$$\left.\frac{\partial < B >_z}{\partial z}\right|_{z=0} = -\frac{< xB >_0}{< x >_0} \tag{45}$$

In particular, with $B = x^n$ we have

$$\left.\frac{\partial < x^n >_z}{\partial z}\right|_{z=0} = -\frac{< x^{n+1} >_0}{< x >_0} \tag{46}$$

If we come back to the expression of $\left.\frac{\partial q(z_1,z_2)}{\partial z_1}\right|_{z_1=0,z_2=0}$, we obtain

$$\beta = \left.\frac{\partial q(z_1,z_2)}{\partial z_1}\right|_{z_1=0,z_2=0} = \frac{\left[-2\frac{<x^3>_0}{<x>_0} + \frac{<x^4>_0}{<x>_0}\right] 2 < x >_0 + \frac{<x^2>_0}{<x>_0}(4 < x^2 >_0 -2 < x^3 >_0)}{4 < x >_0^2} \tag{47}$$

Replacing in the equation (25) and after some simplifications, we obtain the final expression of $\alpha$

$$\alpha = \left| d\left[\frac{-2 < x^3 >_0 + < x^4 >_0 + \frac{2<x^2>_0^2}{<x>_0} - \frac{<x^3>_0<x^2>_0}{<x>_0}}{2 < x >_0^2}\right] e^{-d\frac{2<x^2>_0-<x^3>_0}{<x>_0}} \right| \tag{48}$$

which can also be written as follows

$$\boxed{\alpha = \left|d\beta e^{-dq}\right|} \tag{49}$$

### 3 Summary statistics

Based on these analytical developments, we can now obtain analytical expressions for the summary statistics.

#### 3.1 Diversity : D

The *Prdm9* diversity, written $D$, is defined as

$$D = \left(\sum_i f_i^2\right)^{-1} \tag{50}$$

We now need to calculate $D^{-1} = \sum_i f_i^2$. For that, we can use again the tiling principle :

$$D^{-1} \approx \frac{1}{\tau}\int_0^\infty f^2(t)\,dt = \frac{1}{\tau}\int_0^{z(\infty)} f^2(z)\frac{dt}{dz}\,dz = \frac{1}{\tau}\int_0^{z(\infty)} f^2(z)\frac{1}{\rho f(z)}\,dz = \frac{1}{\rho\tau}\int_0^{z(\infty)} f(z)\,dz \tag{51}$$

Replacing f(z) by its expression from equation (12) and integrating over $z$ gives :

$$\frac{1}{\rho\tau}\int_0^{z(\infty)} -\frac{\alpha}{4\rho}z(z-2\bar{z})\,dz = -\frac{\alpha}{4\rho^2\tau}\left[\int_0^{z(\infty)} z^2\,dz - 2\bar{z}\int_0^{z(\infty)} z\,dz\right] - \frac{\alpha}{4\rho^2\tau}\frac{z(\infty)^3}{3} + \frac{\alpha\bar{z}}{2\rho^2\tau}\frac{z(\infty)^2}{2} \tag{52}$$

But we know that $\bar{z} = \frac{\rho\tau}{2}$ and $z(\infty) = \rho\tau$. So if we replace it in the equation (52)

$$-\alpha\left[\frac{\rho^3\tau^3}{12\rho^2\tau} - \frac{\rho^3\tau^3}{8\rho^2\tau}\right] = -\alpha\left[\frac{\rho\tau^2}{12} - \frac{\rho\tau^2}{8}\right] = -\alpha\left[\frac{2\rho\tau^2}{24} - \frac{3\rho\tau^2}{24}\right] \tag{53}$$

So we obtain

$$D^{-1} = \frac{\alpha\rho\tau^2}{24} \tag{54}$$

But, we also know that $\tau = \frac{2}{\mu\alpha\bar{z}}$ and $\bar{z} = \sqrt{\frac{\rho}{\mu\alpha}}$. So then $\tau = \frac{2}{\mu\alpha}\sqrt{\frac{\mu\alpha}{\rho}}$ and $\tau^2 = \frac{4}{\mu\alpha\rho}$. Which leads to the equation

$$D^{-1} = \frac{4\alpha\rho}{24\mu\alpha\rho} = \frac{1}{6\mu} \tag{55}$$

Where $\mu = 4Nu$, so finally we have

$$D^{-1} = \frac{1}{24Nu} \tag{56}$$

And thus

$$\boxed{D = 24Nu} \tag{57}$$

##### 3.2 Mean age : $\bar{z}$

Using equation (49), $\bar{z}$ can be more directly expressed as :

$$\boxed{\bar{z}} = \sqrt{\frac{\rho}{\mu\alpha}} = \sqrt{\frac{Nvd}{2h}\frac{1}{4Nu}\frac{e^{dq}}{d\beta}} = \boxed{\sqrt{\frac{ve^{dq}}{8hu\beta}}} \tag{58}$$

with $q$ given by the equation (42)

##### 3.3 Mean activity : $<\bar{\theta}>$

If we start from the equation (38) replacing $z$ by $\bar{z}$ we have :

$$\overline{\theta_y(z)} = \theta_y(\bar{z}) = e^{-\gamma(y)\bar{z}} \approx 1 + (-\gamma(y)\bar{z}) \approx 1 - \gamma(y)\bar{z} \tag{59}$$

Averaging over the affinity distribution :

$$\boxed{<\theta_y(\bar{z})> \approx 1 - \bar{z}} \tag{60}$$

##### 3.4 Mean probability of symmetrical binding : $\bar{q}$

If we start from the equation (28) replacing $z$ by $\bar{z}$ we have :

$$\boxed{\bar{q}} = q^{het}(\bar{z},\bar{z}) = \frac{2<x^2>_{\bar{z}} - <x^3>_{\bar{z}} + 2<x^2>_{\bar{z}} - <x^3>_{\bar{z}}}{<x>_{\bar{z}} + <x>_{\bar{z}}} = \boxed{\frac{2<x^2>_{\bar{z}} - <x^3>_{\bar{z}}}{<x>_{\bar{z}}}} \tag{61}$$

The moments $<x^m>_{\bar{z}}$ are given by equation (41) an can be evaluated by numerical integration.

##### 3.5 Mean fertility rate : $\bar{w}$

If we start from the equation (24) replacing $z$ by $\bar{z}$ we have :

$$\boxed{\bar{w} = w(\bar{z}, \bar{z}) = 1 - e^{-dq(\bar{z},\bar{z})}} \tag{62}$$

With $\bar{q} = \frac{2<x^2>_{\bar{z}} - <x^3>_{\bar{z}}}{<x>_{\bar{z}}}$.

### 4 Perturbative development accounting for genetic dosage

The evolution of the frequency of an allele in the population as a function of its age has the same general expression as without dosage :

$$\frac{df}{dz} = \frac{1}{\rho}\left(\frac{w^*(z) - \bar{w}}{\bar{w}}\right) \tag{63}$$

However, for the fitness, we now account for the contribution of homozygotes:

$$w^*(z) = f(t)w^{hom}(z) + (1 - f(t))w^{het}(z, \bar{z}) \tag{64}$$

Linearizing in the vicinity of 0 for $z$:

$$w^*(z) = fw^{hom}(z) + (1-f)w^{het}(z,\bar{z}) \approx fw^{hom}(0)(1-\alpha^{hom}z) + (1-f)w^{het}(0,0)(1-\frac{\alpha^{het}}{2}(z+\bar{z})) \tag{65}$$

Our development is perturbative in the sense that it assumes that gene dosage has a weak impact, and this, because homozygotes are assumed to be rare, i.e. because f is small ($f << 1$). Combined with the weak erosion assumption ($\bar{z} << 1$), this means that we can ignore terms of the order of $zf$. Thus, equation (65) simplifies to :

$$w^*(z) \approx fw^{hom}(0) + (1-f)w^{het}(0,0) - w^{het}(0,0)\frac{\alpha^{het}}{2}(z+\bar{z}) \tag{66}$$

Averaging over the population gives the mean fitness:

$$\bar{w} \approx \bar{f}w^{hom}(0) + (1-\bar{f})w^{het}(0,0) - w^{het}(0,0)\frac{\alpha^{het}}{2}(\bar{z}+\bar{z}) \tag{67}$$

Finally :

$$\frac{df}{dz} = \frac{1}{\rho}\left(\frac{w^*(z) - \bar{w}}{\bar{w}}\right) \approx \frac{1}{\rho}\left[\frac{w^{hom}(0) - w^{het}(0,0)}{w^{het}(0,0)}(f-\bar{f}) - \frac{\alpha^{het}}{2}(z-\bar{z})\right] \tag{68}$$

If we express $\sigma(0) = \frac{w^{hom}(0) - w^{het}(0,0)}{w^{het}(0,0)}$, we obtain :

$$\boxed{\frac{df}{dz} \approx \frac{1}{\rho}\left[\sigma(0)(f-\bar{f}) - \frac{\alpha^{het}}{2}(z-\bar{z})\right]} \tag{69}$$

### References

[1] T. Latrille, L. Duret, and N. Lartillot. "The Red Queen model of recombination hot-spot evolution : a theoretical investigation." In: *Philosophical Transactions of the Royal Society B : Biological Sciences* 372 (2017), p. 20160463. DOI: http://dx.doi.org/10.1098/rstb.2016.0463.

[2] H. Kono et al. "Prdm9 Polymorphism Unveils Mouse Evolutionary Tracks". In: *DNA Research* 21.3 (2014), pp. 315–326. DOI: http://dx.doi.org/10.1093/dnares/dst059.

[3] F. Smagulova et al. "The evolutionary turnover of recombination hot spots contributes to speciation in mice." In: *Genes & Development* 30 (2016), pp. 226–280. DOI: http://dx.doi.org/10.1101/gad.270009.115.

### Supplementary Texte S1 : Measure of Prdm9 diversity in Mus musculus sub-species

#### 4.1 Introduction

Kono *et al.* (2014) genotyped the *Prdm9* ZF-encoding exon in 105 individuals, from 4 subspecies of mice (see Table S2 in [2]). In total, they identified 56 alleles encoding different ZF arrays. To analyze *Prdm9* diversity in natural populations, we focused on data from wild-captured mice (total=79 individuals). Kono *et al.* classified *Prdm9* alleles in 12 clusters (Ca, Cb, Cc, Cd, Ce, Da, Db, Dc, Dd, Ma, Mb, t). It should be noted that some of these alleles that belong to a same cluster share a large fraction of their target sites. For instance, *Prdm9* allele C3H (Da1 in [2]) is closely related to B6 (Da2), and these two alleles share ~30% of their targets ([3]). Similarly, PWD (Ma7) and MOL, which both belong to the Ma cluster share about 13% of their targets ([3]). Thus, the total *Prdm9* allelic diversity probably over-estimates the functional diversity (in terms of *Prdm9* target sites).
We therefore computed two estimates of *Prdm9* diversity:

- Dmax: diversity measured by considering all *Prdm9* alleles individually
- Dmin: diversity measured after grouping of *Prdm9* alleles per cluster

#### 4.2 Summary

| Species | NbIndividuals | NbAlleles | Heterozygous | NbClusters | Dmax | Dmin |
|---|---|---|---|---|---|---|
| M_m_molossinus | 10 | 3 | 20.0% | 2 | 1.80 | 1.60 |
| M_m_domesticus | 20 | 12 | 10.0% | 4 | 6.30 | 2.24 |
| M_m_castaneus | 24 | 23 | 50.0% | 7 | 12.52 | 5.82 |
| M_m_musculus | 25 | 27 | 48.0% | 7 | 18.12 | 2.53 |

The number of individuals genotyped in *M. m. molossinus* is too small to estimate *Prdm9* diversity. In the three other subspecies, the total *Prdm9* diversity ranges from Dmax=6.3 to Dmax=18.12. If we consider the *Prdm9* diversity in terms of clusters of alleles, then diversity ranges from Dmin=2.24 to Dmin=5.82.

The true functional *Prdm9* diversity (in terms of target sites) is most probably between Dmin and Dmax.

### References

[1] T. Latrille, L. Duret, and N. Lartillot. "The Red Queen model of recombination hot-spot evolution : a theoretical investigation." In: *Philosophical Transactions of the Royal Society B : Biological Sciences* 372 (2017), p. 20160463. DOI: http://dx.doi.org/10.1098/rstb.2016.0463.

[2] H. Kono et al. "Prdm9 Polymorphism Unveils Mouse Evolutionary Tracks". In: *DNA Research* 21.3 (2014), pp. 315–326. DOI: http://dx.doi.org/10.1093/dnares/dst059.

[3] F. Smagulova et al. "The evolutionary turnover of recombination hot spots contributes to speciation in mice." In: *Genes & Development* 30 (2016), pp. 226–280. DOI: http://dx.doi.org/10.1101/gad.270009.115.

### Supplementary figures

#### Supporting information

**S1 Fig. A simulation trajectory under a polymorphic regime ($u = 5 \times 10^{-4}$, $N = 5 \times 10^{3}$ and $v = 5 \times 10^{-5}$) with same scale as Fig 3.** In all panels, each color corresponds to a different allele. Note that a given color can be reassigned to a new allele later in the simulation. Successive panels represent the variation through time of (A) the frequency of each *PRDM9* allele and its corresponding (B) the proportion of active sites, (C) the mean affinity of active sites, (D) the probability of symmetrical binding and (E) the fertility. The thick line singles out the trajectory of a typical allele.

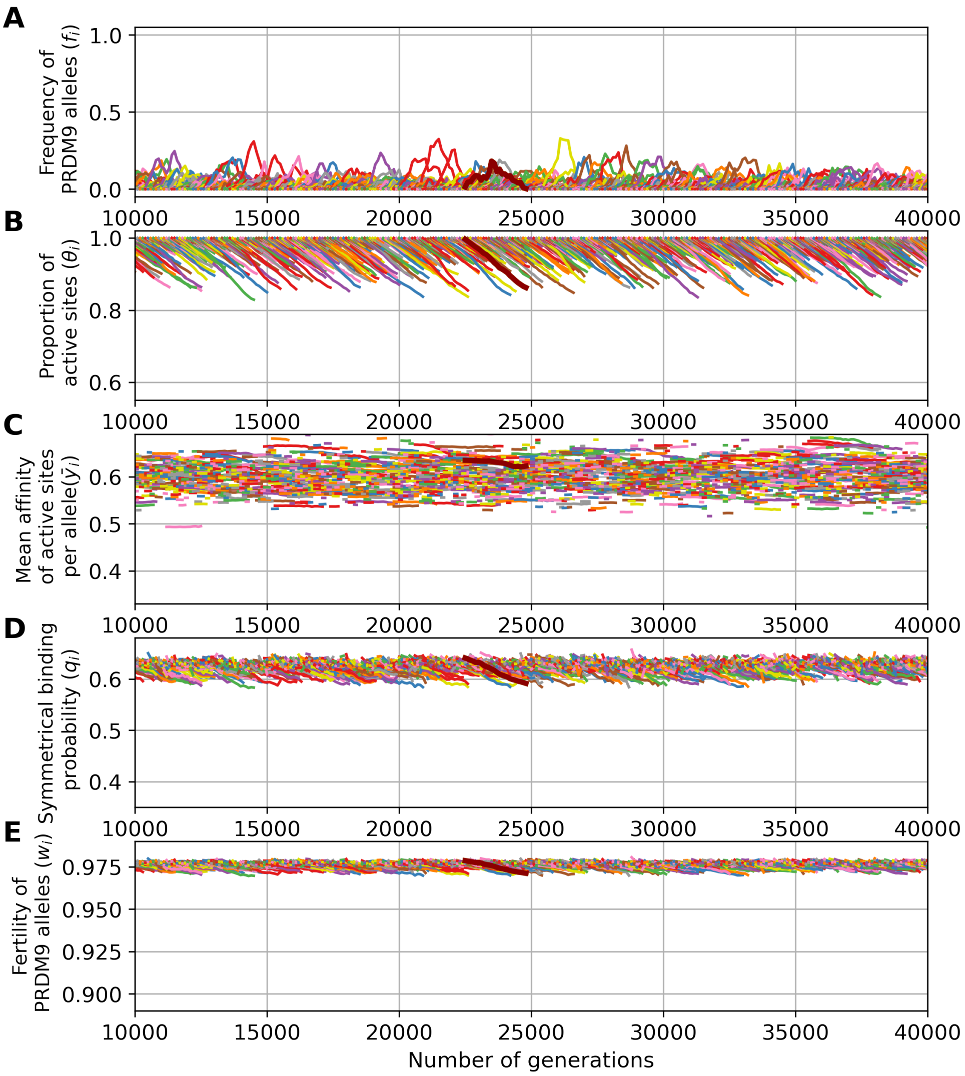

**S2 Fig. Two simulation trajectories under the control model allowing for chromosome pairing and success of meiosis without requiring symmetrical binding of PRDM9.** (A) and (B) correspond to the monomorphic regime (as in Fig 3, $\mathbf{u = 5 \times 10^{-6}}$, $\mathbf{N = 5 \times 10^{3}}$ and $\mathbf{v = 5 \times 10^{-5}}$), while (C) and (D) correspond to the polymorphic regime (as in Fig 4, $\mathbf{u = 5 \times 10^{-4}}$, $\mathbf{N = 5 \times 10^{3}}$ and $\mathbf{v = 5 \times 10^{-5}}$). In all panels, each color corresponds to a different allele. Note that a given color can be reassigned to a new allele later in the simulation. Successive panels represent the variation through time of (A) and (C) the frequency of each *PRDM9* allele and (B) and (D) its corresponding proportion of active sites.

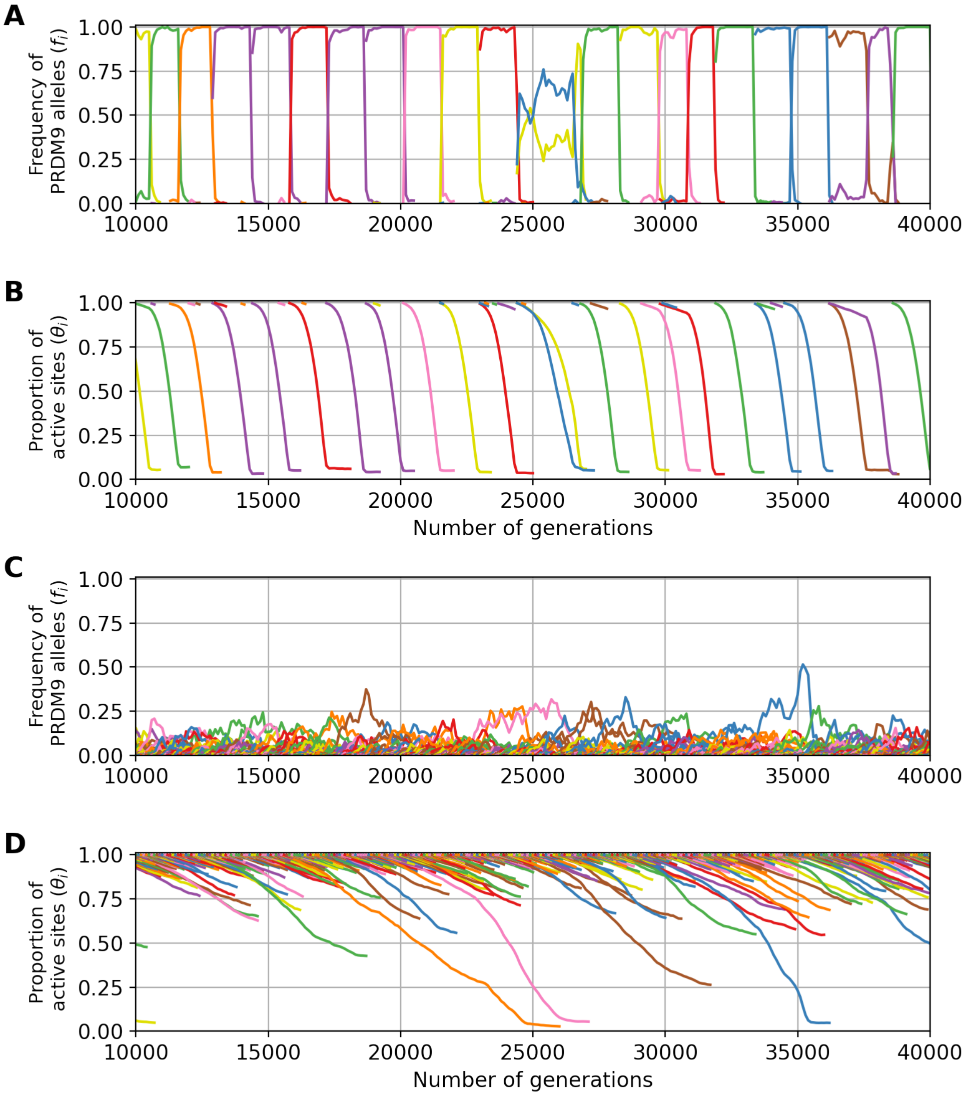

**S3 Fig. A simulation trajectory with genetic dosage and under the control model allowing for chromosome pairing and success of meiosis without requiring symmetrical binding of PRDM9.** The simulation was run with $\mathbf{u = 5 \times 10^{-4}}$, $\mathbf{N = 5 \times 10^{3}}$ and $\mathbf{v = 5 \times 10^{-5}}$. In all panels, each color corresponds to a different allele. Note that a given color can be reassigned to a new allele later in the simulation. Successive panels represent the variation through time of (A) the frequency of each *PRDM9* allele and its corresponding (B) the proportion of active sites.

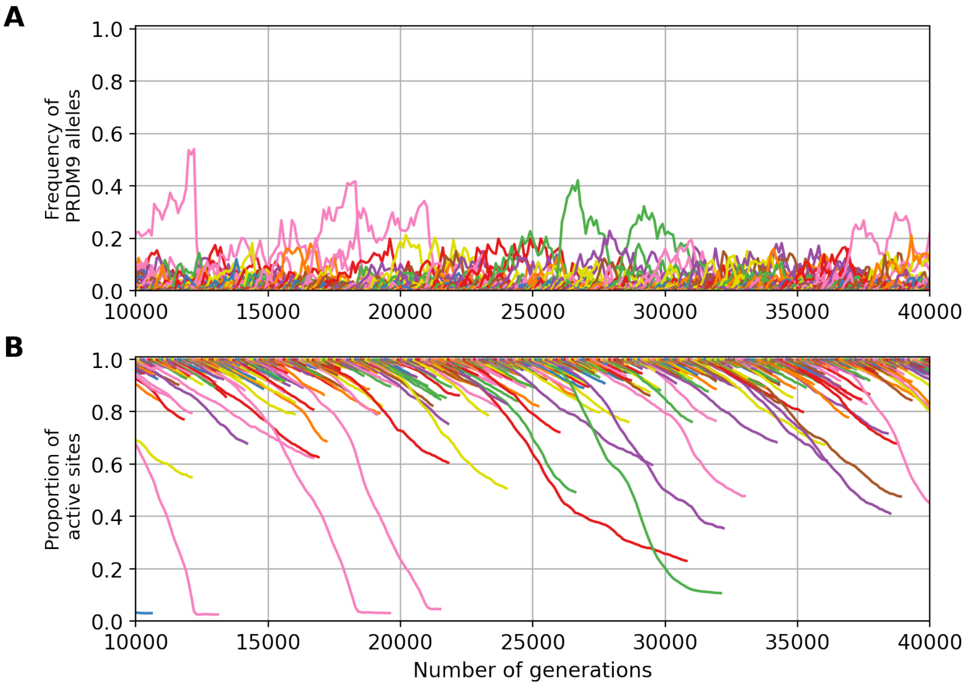

**S4 Fig. Scaling of key summary statistics at equilibrium, as a function of the mutation rate u at the *PRDM9* locus and the mutation rate v at the target sites ($\mathbf{N = 5 \times 10^3}$).** The statistics are: the *PRDM9* diversity $D$ as a function of $u$ (A) and $v$ (D); the mean erosion $\bar{z}$ (i.e. the mean fraction of target sites that have been inactivated) at equilibrium as a function of $u$ (B) and $v$ (E); the mean fertility of the population $\bar{w}$ as a function of $u$ (C) and $v$ (F). On each graph, the mean (blue line) and standard variation (blue vertical bars) over a simulation are displayed against the prediction of the analytical approximation (orange line). The area colored in green corresponds to the range of parameters for which the analytical model verifies the assumptions of a high diversity ($1 < 4Nu < 100$), a low erosion ($\bar{z} < 0.5$) and strong selection on new *PRDM9* alleles ($4Ns_0 > 3$).

The analytical approximations presented here are plotted on panels A to F (orange curves), against the results obtained directly using the simulation program (blue curves). The model and the analytical approximation give qualitatively similar results in the range of parameters validating all the conditions (in practice, we consider that the analytical results should be valid in the following intervals for the model parameters: $1 < 4Nu < 100$, $\bar{z} < 0.5$ and $4Ns_0 > 3$). Concerning *PRDM9* diversity, substantial differences are observed between the simulation results and the analytical approximations, up to a factor of 10, in the scaling of $u$ (panel A). However, the nature of the regime, polymorphic (many alleles segregating at the same time in the population) or monomorphic (only one allele present in the population at a time), is correctly predicted. In particular, we can say that the nature of the regime is directly and mostly determined by $Nu$ and the level of erosion has almost no influence (panel D). Finally, the analytical approximations are less accurate for low and high $u$ or high $v$. These correspond to strong erosion regimes (low $u$ and high $v$) or to regimes with weak selection (high $u$), for which the assumption of the analytical developments are not met.

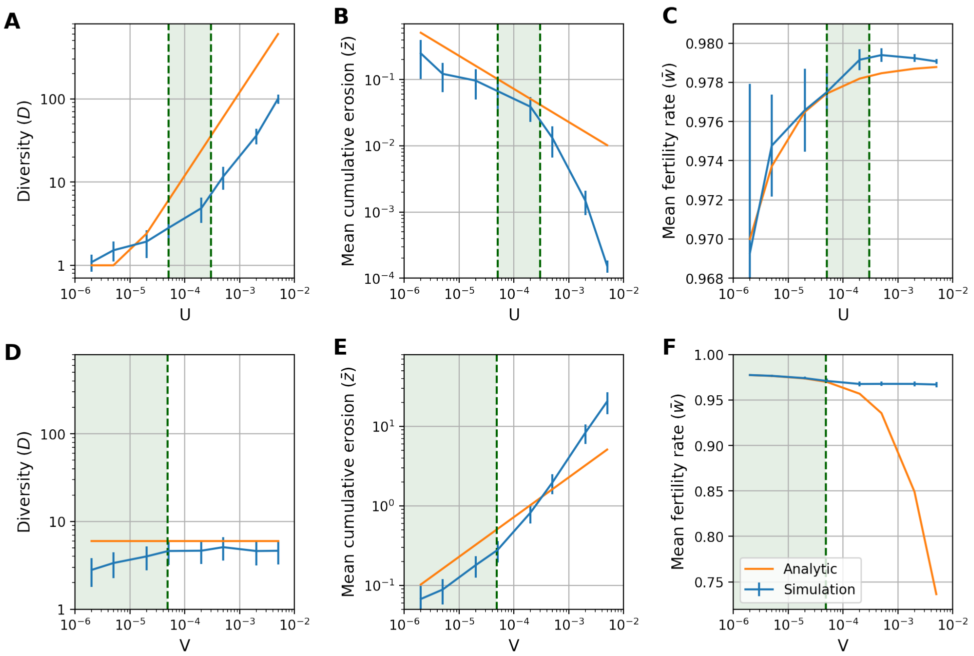

**S5 Fig. Exponential law for affinity distribution (with mean $y = 0.2$).** The continuous blue line corresponds to the affinity distribution for young alleles and the dotted red line corresponds to the affinity distribution for old alleles.

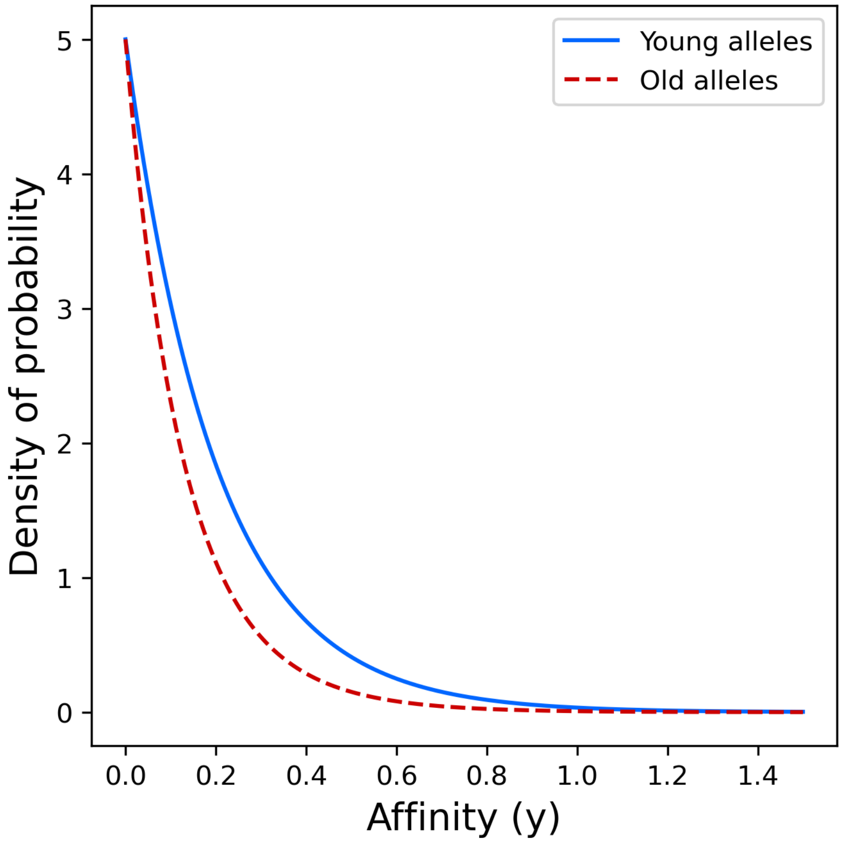

**S6 Fig. Binding probability as a function of the concentration of free PRDM9 protein molecules for homozygotes and heterozygotes (mean affinity $y = 0.6$).** The binding probability for homozygotes is always higher than that for heterozygotes, but the difference between them decreases when the free PRDM9 concentration increases.

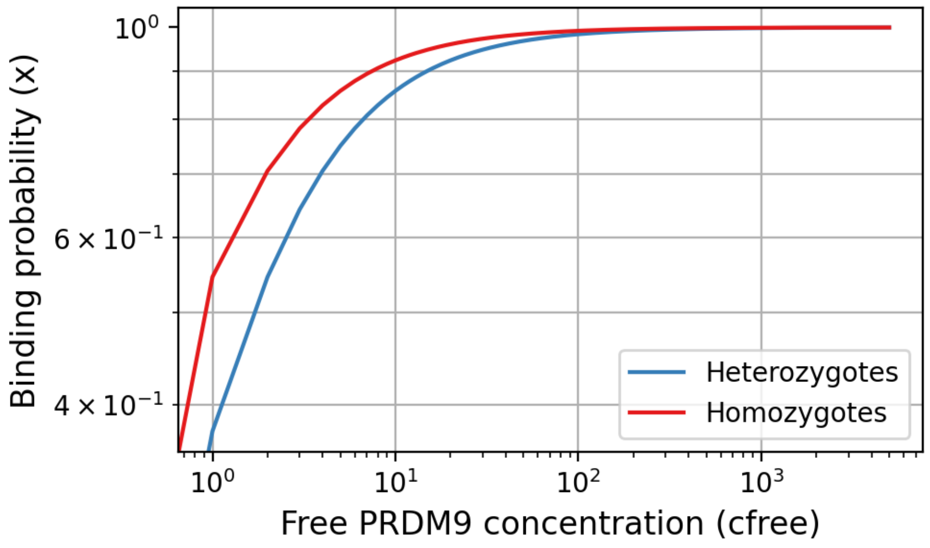


**S7 Fig. Selection coefficient associated to gene dosage ($\sigma_0$) as a function of the concentration of PRDM9 in the cell and the mean affinity of the target sites ($\bar{y}$).**

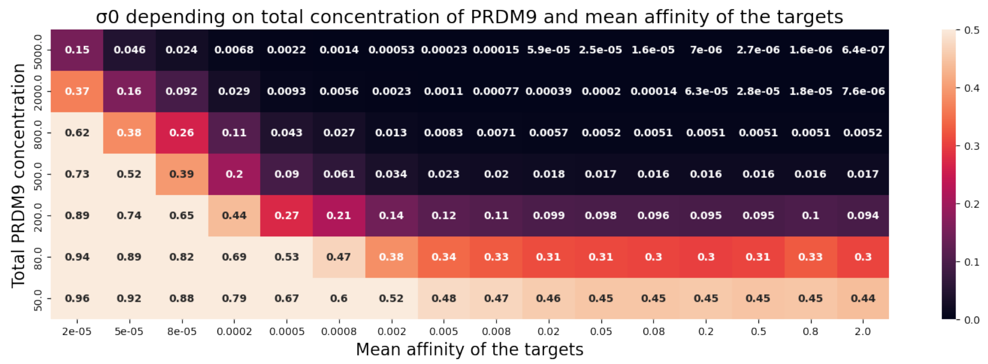
